# Controlled intramural fluid injection to quantify propensity to thoracic aortic dissection

**DOI:** 10.64898/2026.05.10.721415

**Authors:** C. Cavinato, B. Pierrat, E. Ban, M. Scheel, M. Simon, J.D. Humphrey

## Abstract

Dissection of the thoracic aorta includes delamination of medial lamellae and permeation of blood within the media. Quantifying how biaxial loading of a vulnerable wall and fluid mechanics interact to drive dissection remains a central challenge. Here we combine controlled distension-extension testing of intact porcine descending thoracic aortas with forced intramural fluid injection to investigate how axial stretch, injection rate, and needle gauge modulate the initiation and propagation of intramural delamination. Across experiments, injection pressure–volume curves exhibited nonlinear responses characterized by pressure peaks followed by stepwise pressure drops, suggesting progressive micro-delamination events within the medial lamellar networks. Increasing axial stretch significantly elevated peak injection pressure and promoted preferential axial propagation of the permeation / delamination front. Higher injection rates induced abrupt lamellar separation and larger dissected areas, whereas smaller needle gauges generated higher upstream pressures due to increased hydraulic resistance. Synchrotron imaging revealed the microstructural transition from intralamellar fluid permeation and wall swelling to the formation of a large fluid-filled delamination cavity. These results support a mechanistic framework in which the introduction of pressurized fluid within the aortic media behaves as a hydraulic fracture process in a layered poroelastic tissue, governed by balance across fluid pressurization, wall loading, and interlamellar strength. The findings provide quantitative insight into the biomechanical conditions that contribute to the initiation and propagation of aortic dissection.

## Introduction

Thoracic aortic dissection remains a catastrophic cardiovascular event and leading cause of morbidity and mortality [1]. Clinical outcomes depend strongly on the site and extent of the dissection, with proximal dissections (Stanford A) typically treated via emergency surgery and distal dissections (Stanford B) often managed medically by blood pressure control [2, 3]. At the structural level, delamination of adjacent medial elastic lamellae is considered a key mechanism driving aortic dissection. Degenerative changes such as elastin fragmentation and local accumulations of glycosaminoglycans/proteoglycans can reduce interlamellar cohesion and promote intramural separation under mechanical stress or localized pressurization [4, 5].

This failure mechanism is also closely related to aortic intramural hematoma (IMH), an acute aortic syndrome defined by hemorrhage within the media without a visible intimal tear, accounting for 10-30% of cases [6]. Whereas the “inside-out” theory assumes that dissections initiate from an intimal tear, the “outside-in” hypothesis proposed by Krukenberg in 1920 [7, 8] suggests that many dissections originate from IMHs caused by rupture of the vasa vasorum or microscopic intimal defects. Unlike classic dissection, IMH often lacks a primary re-entry tear to allow mechanical decompression, creating a confined pressurized fluid inclusion within the lamellar architecture that generates stress concentrations and promotes rapid delamination. Clinically, 28-47% of IMHs cases progress to overt dissection with a visible intimal flap [9]. Despite its clinical significance, isolating the biomechanical mechanisms that govern the transition from a stable intramural inclusion to a propagating dissection is particularly challenging because the process unfolds within the medial layer under in vivo conditions that are difficult to reproduce or observe [10, 11].

Among the early approaches devised to address these questions was injection of a physiologic solution within the medial layer of the aortic wall, championed by Margot Roach and colleagues [12–14]. This method can provoke lamellar delamination and quantify both the pressure required to initiate intramural tearing and the conditions needed to drive propagation. Early work by Prokop and colleagues, followed by van Baardwijk and Roach [15, 16], further demonstrated the importance of hemodynamic factors that include rate of pressurization (*dP*/*dt*) and pulse pressure *PP*. Carson and Roach [14] found that initiation of a medial tear within a healthy porcine aorta requires supraphysiological pressures while propagation occurs at pressures within the physiological range, thereby distinguishing mechanical thresholds for initiation versus progression. The associated tearing pressure was ∼72 kPa (540 mmHg). Subsequent work explored how local tissue properties, anatomical location, or pathological changes affect medial strength and energy release rate. Roach and Song [13, 17] demonstrated regional differences in porcine thoracic versus abdominal aorta that correlated with microstructural variations in elastin while Tiessen and Roach [13, 17] examined human aortas and highlighted the influence of sex, site, and atherosclerosis. Tam et al. [18] adapted the method to pressurized aortas, revealing that the depth of the initial tear strongly modulated the pressure required for further propagation, with most severe dissections appearing to be in the outer third of the wall. Collectively, these findings suggest that dissection depends on the interplay of global mechanical loads, local wall structure, and tear morphology.

Despite these advances, most injection-based studies have been performed in opened or unpressurized vessels, limiting their physiological relevance. *In vivo*, the aorta is subjected to complex biaxial loading due to luminal pressure and axial tethering, which strongly influence wall mechanics and failure patterns. Moreover, parameters governing the injection, including needle size, insertion depth, and injection rate, have not been systematically characterized even though these parameters determine the local stress concentration and the extent of tissue disruption. These parameters directly control the magnitude and spatial distribution of local intramural stresses and may therefore strongly influence both the onset of lamellar delamination and the subsequent propagation of dissection.

The present study addresses these limitations by combining injection of a physiologic solution during controlled biaxial mechanical loading of the intact porcine descending thoracic aorta (DTA). Using a custom experimental platform, we examined how axial stretch, injection rate, and needle gauge affect injection pressure needed to initiate and propagation a medial delamination as well as histo-morphological outcomes under physiologically relevant conditions. In addition, *in-situ* synchrotron phase-contrast microtomography was used to visualize the three-dimensional microstructural organization of the aortic wall and the morphology of intramural fluid propagation at micron resolution under different injection regimes. By integrating mechanical testing, optical tracking, s-CT imaging, and histological analysis, this work provides novel quantitative insight into the interplay between thoracic aortic loading and intramural flow parameters in the initiation and propagation of aortic dissection.

## Methods

### Preparation of porcine samples and mechanical characterization

Proximal sections of healthy DTAs of standard length (56 ± 3 mm, Figure 1a) were harvested from adult male Yorkshire pigs (25 ± 2 kg) in accordance with protocols approved by the Institutional Animal Care and Use Committee of Yale University. Perivascular tissues were removed and all branches were ligated using nylon sutures. Each vessel was secured on custom-designed ABS cannulas with ligatures and placed in a computer-controlled biaxial distension-extension system [19]. The setup included a rectangular plexiglass chamber filled with a Hanks’ Balanced Salt Solution (HBSS) maintained at room temperature throughout testing, which reduces smooth muscle cell contraction. Pressure and force were monitored using standard transducers, vessel diameter was measured using a video-based optical system, and axial length was prescribed via a stepper motor (Figure 2a).

**Figure 1.**
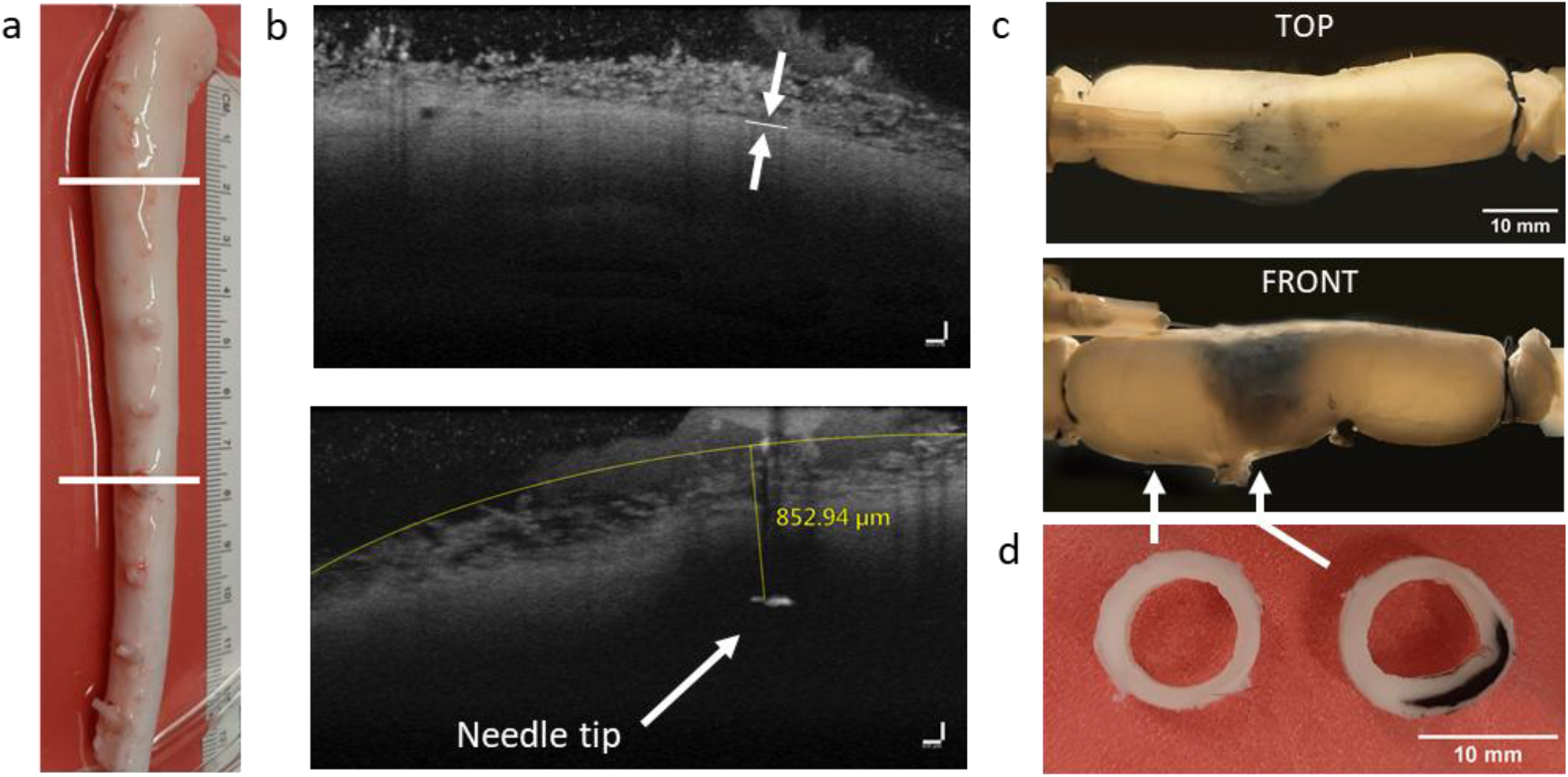
Porcine aortic injection. (a) Proximal section, between the white lines, of the descending thoracic aorta (DTA) used for injection tests. (b) Cross-sectional OCT images of the wall ∼10 mm from the needle tip (top) and at the needle tip location (bottom). Images verify the needle position relative to the adventitia/media interface (marked in the top image, the adventitia being in the upper portion) and measurement of the needle tip depth transverse to the vessel surface (labeled in the bottom image). Scale bar: 100 µm. (c) Camera views from above and lateral to the injection site, with optical axes arranged orthogonally, corresponding to the maximum pressure reached during dissection. Scale bar: 10 mm. (d) Cross-sections taken at the injection site and ∼10 mm upstream from the ink mark. Scale bar: 10 mm.

**Figure 2.**
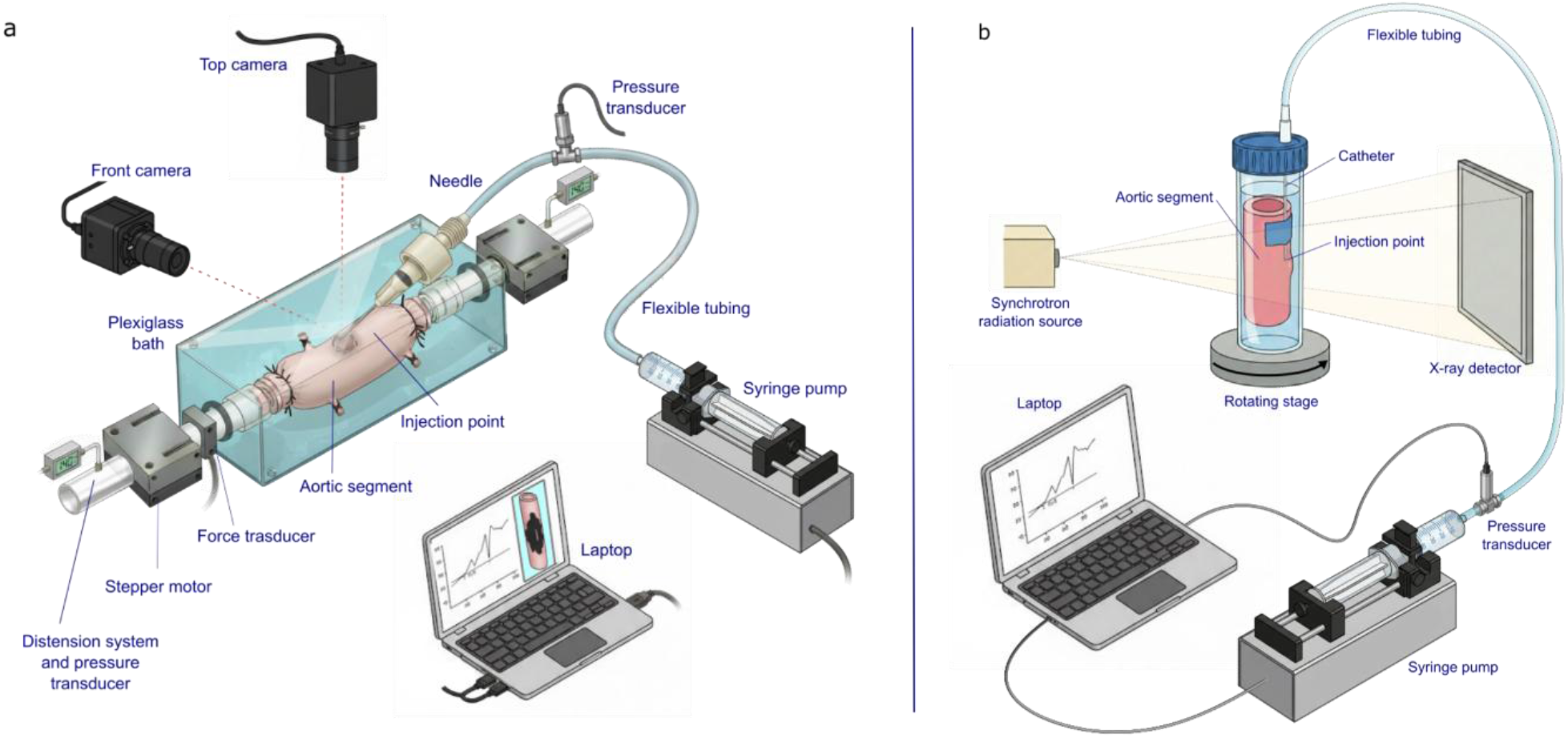
Experimental setups for biaxial distension-extension and intramural injection experiments. (a) Biaxial distension-extension system used for mechanical testing and intramural injection. The aortic segment is mounted on tubes in an HBSS-filled plexiglass bath with axial stretch prescribed by a stepper motor and luminal pressure and axial force monitored on-line by standard transducers. India Ink is injected under pressure via an external needle connected to a syringe pump with upstream pressure measurement, while two orthogonal cameras track the propagation of the ink-filled permeation / delamination front. (b) Synchrotron phase-contrast microtomography setup. The aortic segment containing the catheter injection site is immersed in PBS in a Falcon tube mounted on a rotating stage and imaged between the synchrotron source and the X-ray detector while injection pressure and flow rate are controlled by a syringe pump.

Samples were subjected to equilibration at 90 mmHg for 15 min, then five preconditioning pressurization cycles between 10 and 130 mmHg at a fixed axial stretch of 1.1, with verification of absence of leakage. A randomly selected subset of samples (n = 6) was subjected to biaxial distension-extension mechanical testing. Subsequently, all samples underwent intramural injection experiments. The biaxial protocol consisted of seven loading protocols: (i) three pressure–diameter tests (from 10 to 140 mmHg) at axial *in vivo* stretch (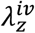, corresponding to the *in vivo* axial stretch at which axial force remains nearly constant with pressure) and ±5% of 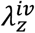; (ii) four axial extension tests at fixed luminal pressures of 10, 60, 100, and 140 mmHg, extended from 0 to *f*_*max*_. Here, *f*_*max*_ is the specimen-specific axial force reached at 140 mmHg and 1.05 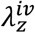.

The unloading portions of the final cycles from all seven protocols were fit simultaneously with a validated four-fiber family constitutive model using Marquardt–Levenberg nonlinear regression [20]. Both the model parameters and the elastic energy stored were determined from these fits, while dissipated energy between loading and unloading curves was also extracted. The stored energy function was:

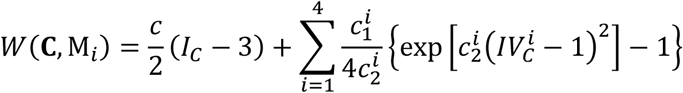

where *c* (kPa), 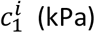, and 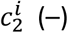 are material parameters. *I*_*C*_ = tr(**C**) and 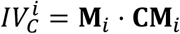 are invariants of the right Cauchy–Green tensor **C** = **F**^*T*^**F**, with **F** = diag[λ_*r*_, λ_θ_, λ_*z*_] and det **F** = 1 (incompressibility). Fiber orientations were defined as axial 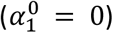, circumferential 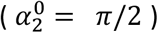, and symmetric diagonal 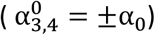.

### Intramural injection experiments

Each vessel was stretched axially to a fixed stretch ratio (1.1 or 1.3) and pressurized to 130 mmHg, equivalent to an elevated physiological systolic arterial pressure in pigs. Samples were randomly assigned to multiple groups: (i) axial stretch ratio of 1.1 vs. 1.3, (ii) injection rate of 1 ml/min vs. 2 ml/min, and (iii) injection needle size of 25G (inner diameter 0.26 mm) vs. 30G (inner diameter 0.16 mm). The injections were performed from the adventitial side using a blunt 1-inch needle (SAI infusion technologies), inserted nearly longitudinally into the vessel wall at the shallowest angle permitted by the surface. The needle tip was positioned within the medial layer, with a penetration depth of ∼1.0 mm for 30G needles and ∼1.5 mm for 25G needles, controlled by a mark on the exposed shaft. For 25G needles, a small backward retraction of ∼500 µm was applied after insertion to enlarge the initial damaged volume. The actual penetration depth of the needle tip was confirmed by cross-sectional acquisitions with Optical Coherence Tomography (SD-OCT Ganymede Thorlabs, resolution of 2 µm over a 2.0 mm imaging depth, Figure 1b) at the needle extremity when penetration allowed, and by side-view camera measurements based on the protruding needle length and insertion angle relative to the vessel surface (Figure 1d). After removal of residual air, a syringe pump delivered HBSS with fixed concentration of India ink at the defined injection rate. The pump was connected to the needle with 2.79 mm diameter tubing (Cole-Parmer®). Injection pressure was measured by a pressure transducer positioned 30 mm upstream of the needle. The injection was continued until observing an evident decrease in injection pressure. To track permeation / delamination propagation, two industrial-grade cameras (Sony IMX179 CMOS sensors, 3264 × 2448 pixels, 1.4 µm pixel size) with 5–50 mm varifocal lenses were mounted around the vessel circumference, above and lateral to the injection site, with their optical axes arranged orthogonally, i.e., 90 degrees (Figure 1c). This configuration provided overlapping yet independent views of the vessel surface recorded at 1 Hz. Focal length was adjusted to ensure adequate magnification and depth of field for accurate reconstruction and registration of the vessel’s external ink-filled permeation / delamination front.

### Correction for tubing and needle effects

Tubing compliance and needle resistance were quantified to correct injection measurements. The system, including a blocked needle held in a metallic needle holder with a ferrule pressure screw, was tested with repeated injections (3 trials for each condition: 25G or 30G, 1 or 2 ml/min). Due to the blocked needle, the measured displacement reflected tubing deformation and pump resistance. Injection curves were nonlinear and fitted by a one-phase association model, *Y* = *Y*_*Plateau*_ · (1 − *e*^−*Kx*^) providing parameters used for volume correction (Supplemental Table S1). Pressure corrections, accounting for needle gauge and hydraulic resistance, were estimated with the Hagen– Poiseuille equation Δ*P* = *Q* · 8μ*L*⁄π*r*^4^, where Δ*P* is the pressure drop [Pa], *Q* the volumetric flow rate [m^3^/s], μ the fluid viscosity [Pa·s], *L* the segment length (tube or needle) [m], and *r* the internal radius [m]. This equation was applied separately to tubing segment and needle resistances. At 1 mL/min, Δ*P* was 36.3 mmHg and 259.2 mmHg for 25G and 30G, respectively; at 2 ml/min, pressure drops doubled (Supplemental Table S2).

### Synchrotron phase contrast microtomography and intramural injection

One proximal segment of porcine DTA obtained as described above, but frozen, was thawed and prepared for synchrotron imaging experiments. A 18G catheter was used for intramural injection to avoid imaging artifacts associated with metallic needles during X-ray acquisition. A pilot channel was first created by laterally piercing the arterial wall with a smaller needle up to approximately mid-wall depth. The catheter was then inserted into this channel and secured using cyanoacrylate glue applied away from the insertion site (Figure 2b). The sample was then immersed in phosphate-buffered saline (PBS) within a Falcon tube and secured to a rotating stage. A syringe pump equipped with a pressure sensor was used to control injection volume and monitor injection pressure. The injected solution consisted of 10% Iomeron-400 (iodinated contrast agent) and 90% PBS, providing radiographic contrast while maintaining near-physiological osmolality [21].

To investigate the microstructural mechanisms at play during two different injection regimes (low vs. high rates), the injection protocol consisted of two steps. A baseline scan was first acquired prior to injection (referenced as “*before injection*”). A slow injection was then performed at 2 mL/min for 50 s, followed by a 10-min stabilization period before acquiring a second scan (“*after slow injection*”). Subsequently, a higher-rate injection was performed at 50 mL/min until delamination. After a 15-min stabilization period, a third scan was acquired (“*after dissection*”). Finally, the injected fluid was withdrawn until pressure returned to 0 kPa, and a final scan was obtained (“*unpressurized*”). For post-processing, injection pressure measurements were corrected for pressure drop along the tubing and catheter based on their diameters and the input injection rates.

X-ray phase contrast microtomography was performed on the ANATOMIX beamline at the SOLEIL Synchrotron, France (Proposal Number 20241557). A volume of 13×13×13 mm^3^ centered at the catheter tip was imaged at a resolution of 2048×2048×2048 voxels (voxel size 6.5 µm), with a scan duration of 2 min 30 s.

### Quantitative histology

Following the injection tests, each vessel was sectioned at the injection site (2 mm thick) to identify the radial position of the false lumen, if any, and an additional section was taken ∼10 mm upstream to obtain a region not damaged by injection (Figure 1d). Traction-free wall thickness was quantified from calibrated camera images. Samples were fixed in 10% formalin for one week, transferred to 70% ethanol for storage, embedded in paraffin, and cut into 4 µm slices. Sections were mounted on slides, heat-fixed, and stained with hematoxylin and eosin. Microscopy was performed with an Olympus BX/51 microscope equipped with a digital camera (4080 × 3072 pixels) and a 10× objective, yielding and effective resolution of 0.21 µm/pixel. Images were captured with Olympus CellSens Dimension software. Given the limited field of view and large samples, multiple images were acquired and stitched using Microsoft Image Composite Editor to reconstruct the arterial cross-section.

### Image reconstruction and analysis

The two orthogonal views of the vessel at the highest injection pressure were segmented to extract the ink-stained regions. Each view was calibrated by matching the measured vessel diameter to its pixel equivalent. Side-view images were co-registered to the top view using a similarity transformation. Morphological filtering and segmentation operations were applied in MATLAB R2025a and segmented contours were then mapped onto cylindrical coordinates, then unwrapped to produce a 2D representation of the vessel surface to facilitate quantification of the permeation / delamination propagation. Morphometric properties were computed with the Matlab tool *regionprops*: the delamination front was approximated by an ellipse, providing major and minor axes, principal with respect to the longitudinal vessel axis, solidity, and circularity. Axial and circumferential extents, as well as area, were quantified to describe the spatial propagation of the lesion.

An anisotropy index was defined along the axis of the cylinder as (*Extent*_*Y*_ − *Extent*_*Z*_)/ (*Extent*_*Y*_ + *Extent*_*Z*_). In this definition, a value of 0 indicates an isotropic spot, values greater than zero correspond to a spot extended along Y (cylinder axis), and values less than zero correspond to a spot extended along X (transverse direction). Solidity was defined as the ratio between the segmented area and its convex hull, with lower values reflecting irregular or fragmented boundaries. Circularity was computed as a ratio of area and perimeter, with values closer to 1 indicating more regular and compact shapes and smaller values corresponding to elongated or irregular profiles.

Synchrotron-CT (s-CT) volumes were segmented into five classes (i.e., catheter, aortic segment, injected fluid, connective tissue, and background) using a 2.5D Attention U-NET with a depth level of 4 and 5 input slices, implemented in Dragonfly 3D World 2025.1 software. Training data consisted of manually segmented slices (∼12 per timestep) augmented ten-fold using geometric and intensity transformations (horizontal/vertical flips, 180° rotation, zoom, shear, brightness). The network was trained using the AdamW optimizer with a cross-entropy loss function for 70 epochs. Segmented fluid regions were subsequently used to quantify local interlamellar defects, delamination morphology, and volumetric distributions associated with intramural fluid propagation.

Histo-morphometric assessment of H&E-stained cross-sections was performed by tracing the boundaries between non-infused lamellae and dark-infused or dissected lamellae to identify lamellar microfractures and local interlamellar decohesion associated with intramural fluid propagation.

### Statistics

All data were analyzed using two-way non parametric Scheirer Ray Hare test to evaluate the effects of axial stretch ratio (1.1 vs. 1.3), injection rate (1 vs. 2 mL/min), and needle size (25G vs. 30G). When significant effects were detected, post hoc Tukey tests were performed. Statistical significance was denoted as follows: \*\**p* < 0.001, \**p* < 0.01, *p* < 0.05, and “ns” for non-significant results.

## Results

### Mechanical characterization

Figure 3 shows mean Cauchy stress–stretch curves for intact healthy porcine DTA specimens (n = 6): cyclic pressure–diameter tests at 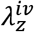 and ±5% of this stretch (panel a) confirmed the typical nonlinear stiffening with increasing pressure as well as modest sensitivity to axial extension while cyclic axial force-length tests at fixed pressures ranging from 10–140 mmHg (panel b) revealed a pressure-dependent stiffening, with specimens elongating up to *f*_*max*_. These results highlight the nonlinear, anisotropic behavior of the healthy proximal porcine DTA. Shown, too, are values of select mechanical metrics and best-fit values of the material parameters used in quantifying the native biaxial properties (Figure 3c, d).

**Figure 3.**
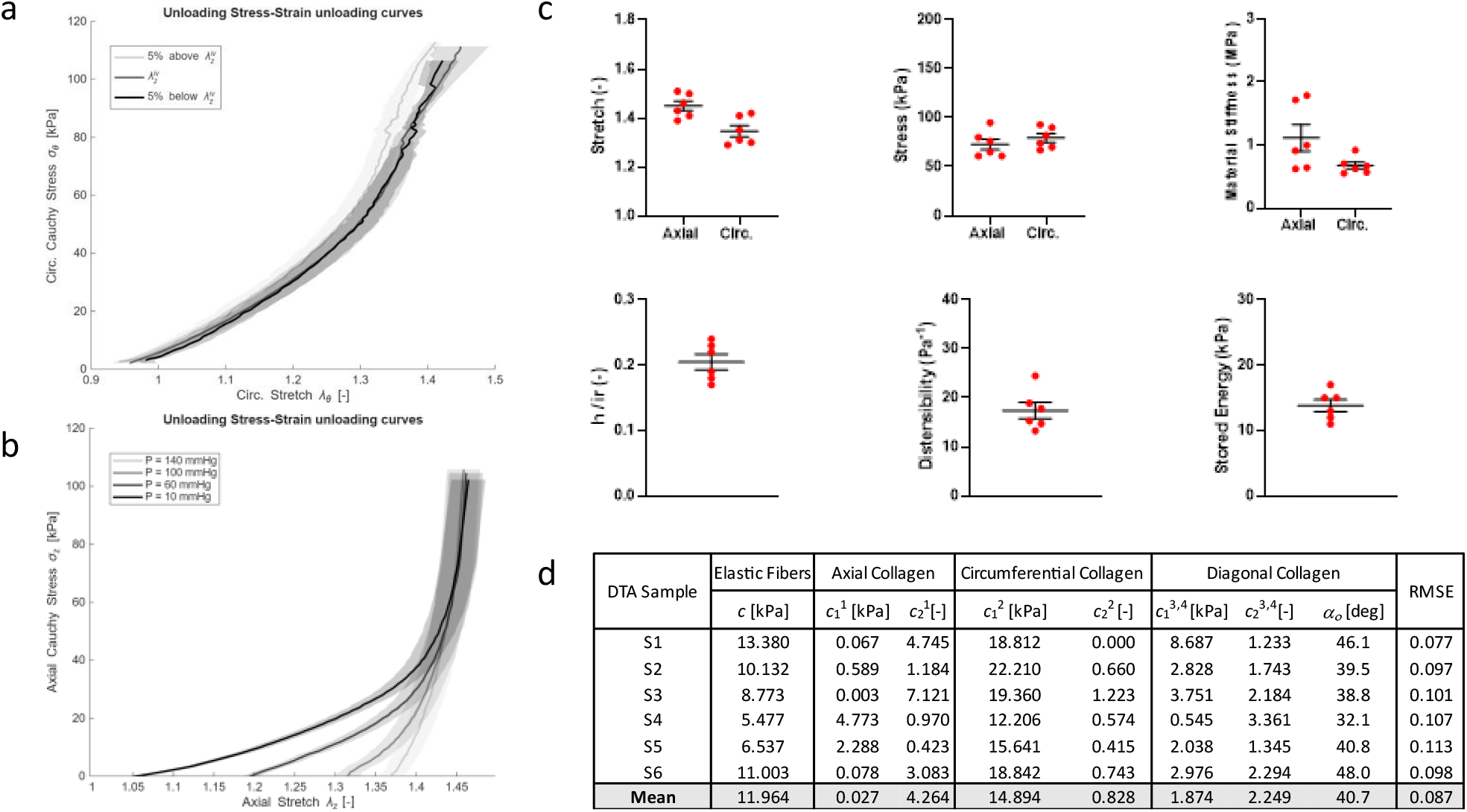
Mechanical behavior of porcine DTA. Mean Cauchy stress–stretch curves (n = 6, average in solid line and SEM in shadow area) in the two principal loading directions, (a) circumferential, obtained from pressure–diameter tests at a fixed in vivo axial stretch (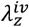, corresponding to the axial stretch at which axial force remains nearly constant during pressure changes) and at ±5% thereof, (b) axial, obtained from axial force-extension tests at fixed luminal pressures of 10, 60, 100, and 140 mmHg, extended up to f_max_. (c) Mechanical descriptors for DTA samples at 120 mmHg and 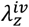s, including stretch, Cauchy stress, and material stiffness in both axial and circumferential directions, stored elastic energy, distensibility between 120 and 80 mmHg (representing typical systolic and diastolic values), and ratio of thickness-to-inner radius at 120 mmHg. (d) Best-fit values of the eight material parameters obtained from optimization of the four-fiber family constitutive model, either for each sample (n = 6) or for the mean experimental curve, with corresponding RMSE values.

### Intramural injection: permeation and delamination characterization

Across all groups, the original pressure–volume injection curves exhibited a characteristic nonlinear profile: pressure rose steeply until reaching a peak, followed by a more gradual, step-like decline. Only in five cases (one sample in the 25G, λ_*z*_= 1.3, 1 mL/min group, two samples in the 30G, λ_*z*_ = 1.1, 1 mL/min group, and two samples in the 25G, λ_*z*_= 1.1, 1 mL/min group) did the curves display an abrupt pressure drop (Figure 4a). For most injections, the step-wise pattern ultimately transitioned into a sudden loss of pressure, typically after several milliliters of fluid had permeated the wall, predominantly into the adventitia. Some experiments were terminated earlier due to substantial leakage of ink into the surrounding bath. Interestingly, the step-wise pattern seen in several curves suggests successive micro-failures or internal ruptures within the media prior to depressurization of the created false lumen. Despite this common pattern, interesting differences were observed among groups on average. Quantification of peak pressures and corresponding injection volumes after correction (cf. Section ***Correction for tubing and needle effects***) confirmed these trends (Figure 4b). Needle penetration depth did not show a clear relationship with either peak injection pressure or injected volume across experimental groups (Figure S1a,b). Injections performed with the smaller 30G needle at λ_*z*_ = 1.1 and 1 mL/min exhibited higher pressures for a given volume compared to the 25G needle. Increasing axial stretch from λ_*z*_ = 1.1 to 1.3 significantly increased the peak injection pressures and injected volumes. Conversely, increasing the injection rate to 2 mL/min shifted the curves upward, with sharp pressure rises occurring at lower injected volumes. These trends were reflected in the summary data even before pressure and volume correction for tubing and needle effects (cf. Figure S2).

**Figure 4.**
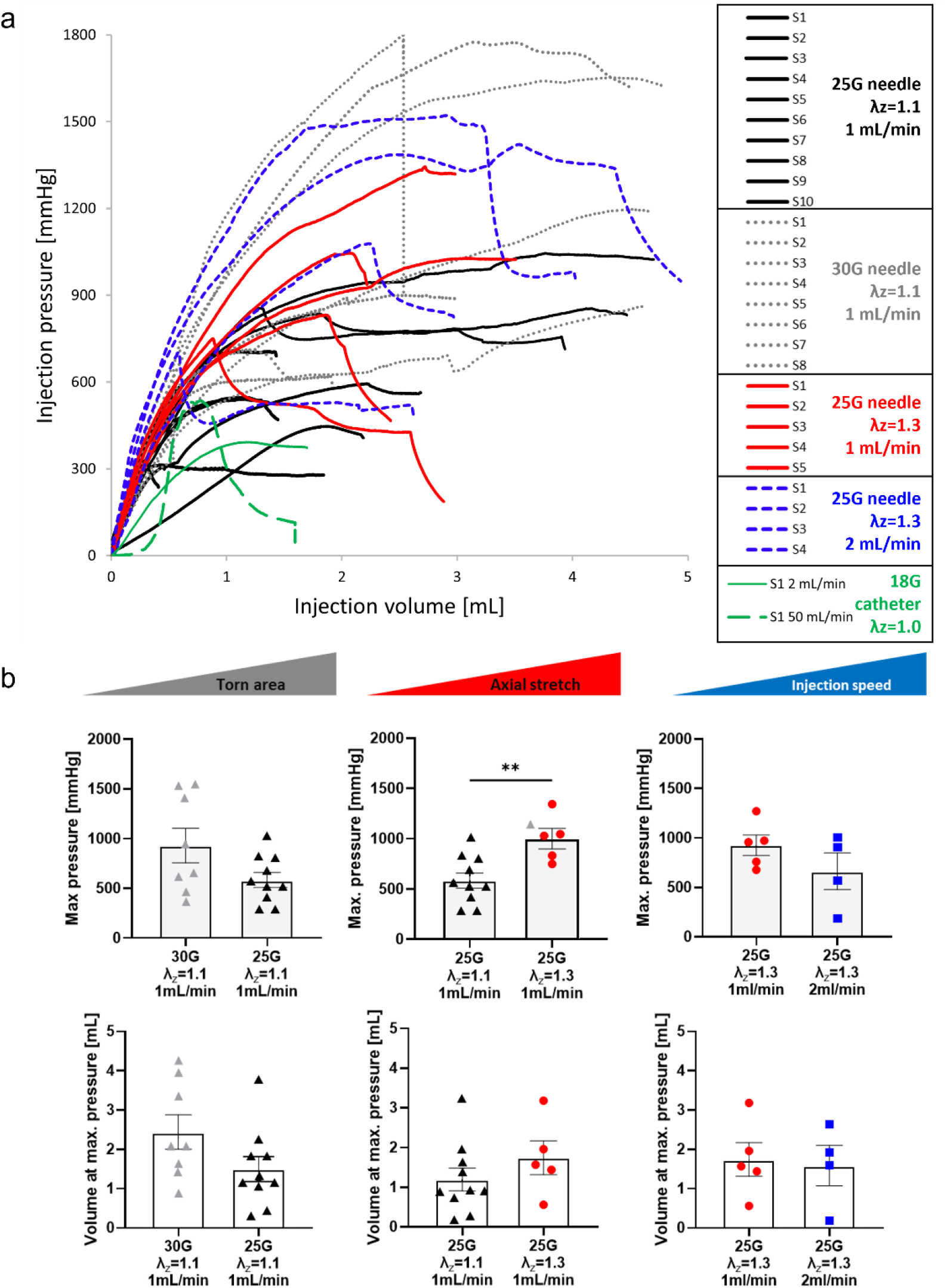
Pressure-volume behavior during injection. (a) Pressure–volume curves for porcine DTAs injected using different experimental conditions in the legend: needle gauges (25G vs.30G), stationary axial stretch during the injection (λ_z_ = 1.1 or 1.3), and injection rates (1 or 2 mL/min), all while fixing the luminal pressure at 120 mmHg. Black solid lines: 25G needle, λ_z_ = 1.1, 1 mL/min; dotted grey lines: 30G needle, λ_z_ = 1.1, 1 mL/min; red solid lines: 25G needle, λ_z_ = 1.3, 1 mL/min; blue dashed lines: 25G needle, λ_z_ = 1.3, 2 mL/min; green solid lines: 18G catheter, λ_z_ = 1.0, 2 mL/min; green dashed lines: 18G catheter, λ_z_ = 1.0, manually imposed 50 mL/min. (b) Bar plots summarizing mean maximum pressures (top) and corresponding injection volumes (bottom) for the four groups after correction for tubing and needle effects (non-corrected, raw data presented in Figure S2). Individual symbol types denote experimental conditions (axial stretch and injection rate) and the x-axis separates the groups according to needle gauge. Error bars represent SEM. Statistical significance, when present, is indicated by horizontal connectors above the bars: **p < 0.001, *p < 0.01, p < 0.05.

Cross-sections of the samples after injection, with corresponding histological overlays, revealed relationships between the injected ink patch and native aortic wall. All four tests performed at 2 mL/min showed an abrupt and extended delamination of the medial lamellae, clearly visible in the cross-sectional images (Figure 5a). Under all other conditions, the lamellar structure was generally preserved except for one sample (25G, λ_*z*_ = 1.3, 1 mL/min) where a tear appeared locally and propagated into the intima. In this case, the injection behaved more as a localized infusion within the wall, rather than evolving into a full delamination. Top-view images (Figure 5b) and cylindrical reconstructions of injections from outer views (Figure 5c) provided a quantitative view of their surface distribution. At low stretch and low flow rate, injected regions remained small and localized whereas the higher stretch and higher injection rate produced larger, elongated stained areas covering a greater portion of the wall. Histological sections further suggested distinct microstructural responses across loading conditions, with localized lamellar decohesion predominating at λ_*z*_= 1.1 and burst-like lamellar delamination observed more consistently at λ_*z*_ = 1.3 (Figure S1c).

**Figure 5.**
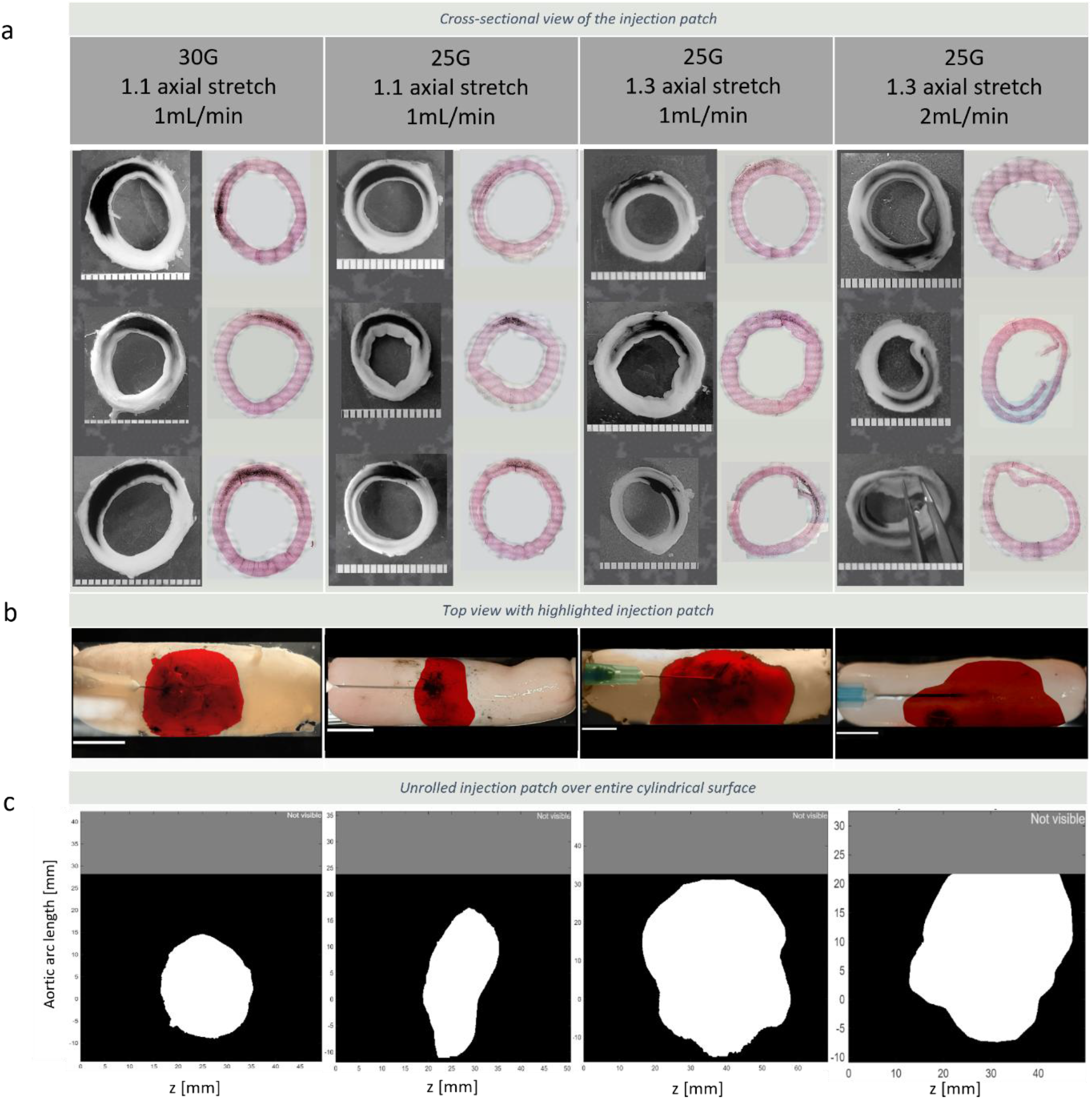
Morphology and distribution of intramural delamination induced by controlled injection under different experimental conditions. (a) Representative cross-sectional views after the injection test (left column) and H&E-stained histological sections (right column) show the location of the permeation / delamination front and interlamellar separation across experimental groups having different values of needle gauge, axial stretch ratio, and injection rate. (b) Top-view images of the aorta with ink-stained regions used to segment the damage front (red area). Scale bar: 10 mm. (c) Cylindrical surface reconstructions obtained by registering orthogonal camera views and projecting vessel contours in cylindrical coordinates (z is the longitudinal length, y-axis is the arc length along the circumference). White regions indicate the effective injection and delamination area within the vessel wall. Morphometric analysis was performed on these segmented and registered images.

Quantitative analysis of the reconstructed injection patches confirmed the qualitative trends (Figure 6), with the delaminated area significantly larger for injections performed at λ_*z*_ = 1.3 compared to λ_*z*_ = 1.1 and the greatest increase observed when the injection rate was raised to 2 mL/min at λ_*z*_ = 1.3. The axial extent of injected volume increased consistently at λ_*z*_ = 1.3 under both flow conditions, suggesting that axial stretch might be a key driver of axial propagation. Additionally, circumferential extent expanded markedly only when the injection rate was raised to 2 mL/min, suggesting that higher flow could promote lateral spreading of the delamination front around the aortic circumference. Anisotropy analysis revealed that patches generated at low axial stretch were more isotropic and compact, while higher axial stretch promoted elongated morphologies. Orientation measurements confirmed this tendency, with injection patches showing a stronger alignment toward the circumferential direction at λ_*z*_ = 1.1 and getting closer to a less defined angle at λ_*z*_ = 1.3. Other shape descriptors provided limited additional insight. Circularity and solidity showed no clear group-dependent differences, while the ratio of vessel enlargement at peak infusion compared to the initial diameter also remained largely unchanged across conditions. Overall, morphometric parameters highlighted the strong influence of both axial stretch and injection rate: higher mechanical axial stretch and faster delivery produced larger, anisotropic delaminations with broader circumferential involvement, whereas low stretch and slower delivery resulted in smaller, more localized patches.

**Figure 6.**
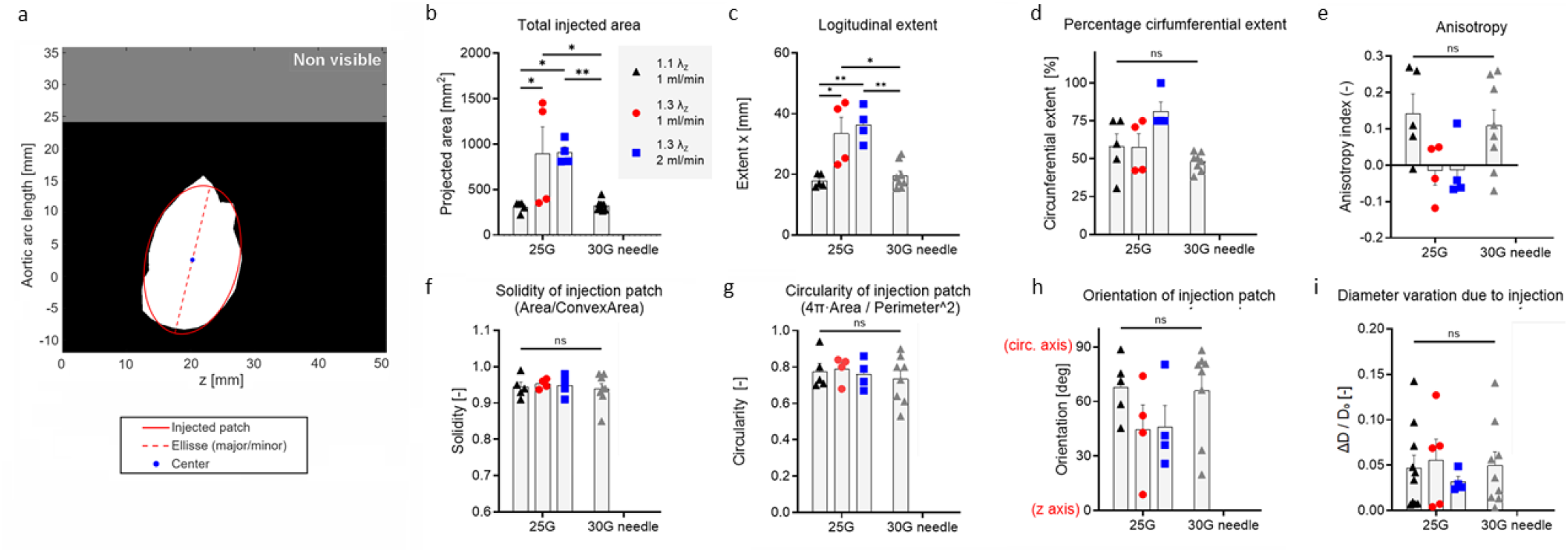
Quantification of injection-induced intramural delamination patches. (a) Representative cylindrical surface reconstructions obtained by registering orthogonal camera views, with z the longitudinal length and y-axis the arc length along the circumference. The white region indicates the segmented injected area within the vessel wall. Morphometric analysis was performed on the mask and the inscribed ellipse. (b–i) Morphometric measurements derived from cylindrical surface reconstructions of the ink-stained wall, including total permeation / delamination area, anisotropy, axial extent (in mm), circumferential extent (in percentage), preferential orientation, circularity, and solidity of the shape. Bar plots show mean values ± SEM for all experimental conditions, with individual data points overlaid (triangle: 1.1 axial stretch, 1 mL/min; circle: 1.3 axial stretch, 1 mL/min; square: 1.3 axial stretch, 2 mL/min). Horizontal connectors indicate statistically significant differences between groups at **p < 0.001, *p < 0.01, p < 0.05.

### Synchrotron phase contrast microtomography and intramural injection

The segmentation model applied to s-CT volumes achieved a loss of 0.0148 and a Dice score of 0.990, indicating excellent segmentation accuracy without the need for further manual refinement. The s-CT acquisitions before injection reveal the lamellar architecture of the arterial tissue (Figure 7a - *before injection*). The initial damage caused by the needle and catheter insertion is visible as delaminated and sectioned lamellae, alongside a small region near the catheter tip where the contrast agent permeated the tissue. Because the artery was not pressurized, the lamellae appeared slightly undulated.

**Figure 7.**
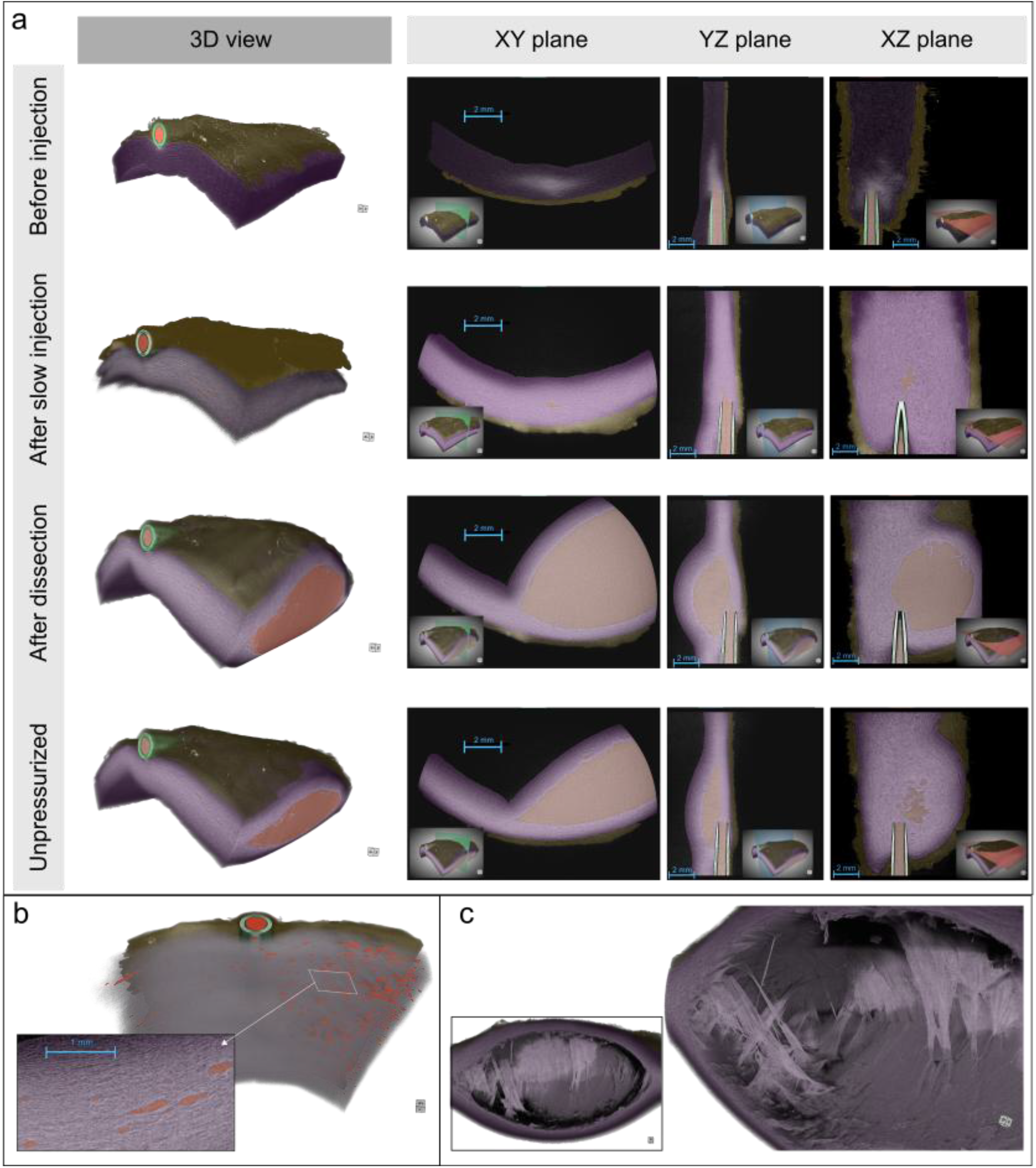
Synchrotron phase-contrast microtomography of intramural injection and delamination in porcine aorta at two different injection rates. (a) Three-dimensional renderings and orthogonal slices (XY and YZ planes) of the arterial wall at four stages of the experiment: before injection, after slow injection, after delamination with high-rate injection, and after depressurization while high-rate injection induces a large interlamellar delamination cavity. (b) Segmentation of localized fluid pockets associated with micro-damage and lamellar decohesion near the catheter tip during the fluid permeation and swelling of the wall under slow injection. The inset highlights representative defects identified as belonging to the injected fluid class. (c) Magnification of the cross-section of the delamination cavity, showing the irregular morphology of the separated medial lamellae and the contrast-agent-filled false lumen.

During the slow injection step, the pressure increased to 55 kPa before plateauing slightly below this value. After the injection stopped, the pressure gradually relaxed to 5 kPa. The contrast agent penetrated deep into the tissue (Figure 7a - *after slow injection*), and the aortic thickness in the region of interest increased from 2.0 to 2.5 mm (approximately 25%) due to swelling. This penetration caused additional micro-damage to the medial lamellae, evidenced by local decohesion in several spots near the catheter. The segmentation successfully identified these local defects as belonging to the fluid class, enabling visualization of their distribution within the arterial wall (Figure 7b). Their volumes followed a log-normal distribution with µ = -7.8 and σ = 1.13, corresponding to a median volume of approximately 4 × 10^−4^ mm^3^.

The delamination event at high-rate injection was characterized by a sudden bulging of the artery (Figure 7a - *after dissection*). The pressure increased to 90 kPa before dropping abruptly. The s-CT images revealed a large delamination cavity filled with contrast agent. At this stage, the lamellae were nearly straight, losing their initial undulation, likely due to the tensile stress induced by the internal injection pressure. Delamination occurred between medial lamellae, closer to the adventitia than to the inner media, near the catheter tip. The delamination profile, visualized in Figure 7c, exhibited a complex and chaotic microstructure with partial delamination of multiple lamellae. After depressurization, the lamellae recovered their undulated shape, although permanent inelastic deformations and damage such as delamination and local decohesion remained.

## Discussion

Intramural injection experiments performed under controlled biaxial loading provide a more *in vivo* relevant view of the mechanisms governing fluid permeation and delamination within the aortic wall. The camera-tracking experiments identified consistent trends across multiple experimental conditions — namely axial stretch, needle gauge, and volumetric flow rate — while the phase-contrast s-CT experiment provided detailed three-dimensional insight into local interactions between fluid pressure and the lamellar microstructure. Although the experimental configurations differ, the two approaches are complementary: the camera experiments capture macroscopic propagation patterns across many samples whereas s-CT resolves the underlying microstructural processes that control the transition from permeation to delamination.

Across all experimental groups, the pressure–volume curves displayed a consistent nonlinear pattern characterized by a rapid pressure increase followed by a sequence of partial pressure drops before final depressurization. This behavior indicates that intramural injection does not immediately generate a single catastrophic delamination. Rather, it suggests progressive fluid permeation and structural rearrangement within the medial lamellar structure. The step-like pressure decreases are consistent with successive micro-failures or localized separations between lamellae [10, 22, 23], gradually creating preferential pathways for fluid permeation. Such progressive damage mechanisms are supported by histology and by the higher-resolution s-CT observations, which revealed local decohesion events preceding the formation of larger interlamellar separations. This pressure–volume behavior differs from that reported in the classical intramural injection experiments of Roach and colleagues [12, 17], as we show in Figure S3. In their studies on pressurized thoracic aortas, intramural injection of saline through a 25-gauge needle produced a characteristic pressure–volume response consisting of a sharp pressure peak corresponding to the initial tearing event, followed by a relatively stable propagation phase at much lower pressure. Roach and Song nevertheless also reported mixed tearing responses across aortic regions, with abdominal aortic segments exhibiting more progressive pressure–volume responses compared with the sharper tearing responses typically observed in thoracic specimens [13]. In the present experiments, pressure–volume curves were more consistently characterized by progressive pressure buildup with multiple partial pressure decreases prior to final depressurization, illustrating a mixed tearing response. Direct comparison between these studies remains difficult because intramural tearing appears highly sensitive to both medial microstructure and experimental boundary conditions. Previous experimental and computational studies have shown that regional differences in aortic microstructure — particularly the density and organization of interlamellar elastin radial struts — can strongly influence tearing behavior and promote either more progressive or more abrupt delamination responses [22, 24]. In particular, Ban et al. demonstrated through combined experiments, multiphoton microscopy, and phase-field simulations that structurally significant radial struts can locally confine fluid propagation and generate step-wise tearing responses through successive rupture events. In addition, differences in hydraulic boundary conditions— including needle resistance, insertion geometry, axial loading, luminal pressurization, vessel configuration, and tissue preservation—may alter the local balance between fluid permeation, interlamellar decohesion, and delamination propagation. The present observations therefore likely represent a different mechanical manifestation of the same underlying hydraulic fracture process rather than a fundamentally distinct mechanism.

The influence of experimental parameters on the pressure–volume response further supports this interpretation. Injections performed with the smaller needle gauge generated higher pressures for a given injected volume than those performed with the larger needle. This behavior is expected from fluid mechanics: the smaller needle lumen increases hydraulic resistance and requires a higher upstream pressure to sustain the imposed volumetric flow rate. In addition, the smaller needle likely produces a smaller initial mechanically damaged region at insertion, increasing the pressure required to initiate intramural fluid propagation. These results highlight the sensitivity of the early stages of intramural injection to the local hydraulic boundary conditions imposed at the needle tip.

Axial stretch also had a strong influence on both pressure generation and damage propagation. Increasing axial stretch significantly increased peak injection pressures and injected volumes prior to depressurization. This response can be interpreted in light of the anisotropic mechanical behavior of the aortic wall. Axial stretch increases the tensile loading carried by collagen fibers oriented obliquely within the medial lamellae, effectively stiffening the wall and increasing resistance to interlamellar separation [25–28]. Consequently, higher pressures are required to initiate and propagate fluid-filled separations. At the same time, once fluid pathways are created, the imposed axial tension promotes preferential propagation along the vessel axis, consistent with the increased axial extent observed in the morphometric analysis.

Injection rate was another key determinant of intramural damage. Increasing the injection rate shifted the pressure–volume curves upward and led to abrupt lamellar separation in all tested samples. Cross-sectional observations confirmed extensive medial delamination under these conditions, consistent with a transition from diffusion-dominated permeation to pressure-driven fracture. At lower injection rates, fluid pressure can diminish through interstitial permeation within the lamellar structure. At higher rates, however, the rate of pressure buildup exceeds the ability of the tissue to drain fluid through its low permeability, causing pore pressure to rise until the interlamellar strength is exceeded [29]. The resulting separation creates a hydraulic pathway that resembles the delamination process directly observed in the s-CT experiment. These observations are consistent with prior computational studies of intramural tearing [22, 24], which demonstrated that pressure-driven delamination depends not only on hydraulic loading but also on local wall stiffness, tearing energy, interlamellar connectivity, and the evolving geometry of the false lumen.

The morphometric analysis of the reconstructed injection patches further clarifies the mechanical drivers of delamination propagation. At low axial stretch and low injection rate, injected regions remained compact and relatively isotropic, indicating that fluid remained largely confined within localized interlamellar spaces. By contrast, higher axial stretches and faster injection rates produced larger, elongated regions with greater circumferential involvement. This observation suggests that axial tension modifies the microstructure of the media, facilitating propagation along the direction of applied load while limiting radial expansion across lamellae. Circumferential spreading, in contrast, appeared to be controlled primarily by injection rate, with higher flow conditions enabling the delamination front to expand more readily around the vessel circumference. Needle gauge also affect the local fluid mechanics at the injection site. For a given imposed volumetric flow rate, a smaller needle lumen increases hydraulic resistance and upstream pressure while increasing the outlet velocity, which can enhance local shear and initiate interlamellar separation. Conversely, larger needles distribute the injected fluid over a wider area, reducing local velocity gradients.

Direct quantitative comparisons across all experimental groups must nonetheless be interpreted cautiously. The camera-tracked injections involved smaller needles, controlled axial stretch, and lower injection rates; the s-CT experiment employed a larger needle, no imposed axial stretch, and a higher injection rate. Each of these parameters influences the hydraulic boundary conditions and the local mechanical state of the wall. Even small variations in needle size, insertion depth, or loading configuration can significantly affect the pressure required to initiate intramural fluid propagation, as also reported in previous injection studies such as those of Roach and colleagues. Importantly, however, these differences do not represent conflicting observations. Rather, they probe complementary aspects of the same mechanical response.

The s-CT experiment further demonstrates the ability of phase-contrast imaging to directly visualize the internal lamellar architecture during fluid injection. Unlike conventional histology, which requires destructive sectioning, or clinical imaging modalities such as CT or MRI, which lack sufficient spatial resolution, phase-contrast s-CT provides a non-destructive three-dimensional view of the medial structure. This enabled direct observation of the transition from interlamellar permeation to lamellar separation at micrometer resolution.

Taken together, the combined experiments reveal a coherent mechanical picture of intramural fluid propagation. At low hydraulic loading, fluid permeates the interlamellar spaces of the media, producing wall swelling and localized micro-damage within a poroelastic regime. As hydraulic load increases—either through higher injection rates or mechanical conditions that increase the effective resistance—and pore pressure rises until the interlamellar strength is exceeded, this triggers a fracture-like delamination. This transition can be described through a global energy balance in which the power supplied by the syringe pump *PQ*_*in*_ is partitioned between poroelastic dissipation *D*_*poro*_ and the energy required to create new fracture surfaces *G*_*Ic*_ *dA*/*dt*, where *G*_*Ic*_ is the mode I energy release rate and *A* the fracture surface area. In the permeation regime, the input power is dissipated through fluid flow within the porous tissue, whereas once the critical pressure is reached, the stored elastic energy drives the propagation of an interlamellar delamination that forms a path of least hydraulic resistance [30, 31].

Within this framework, a critical injection threshold *Q*_*th*_ = α*P*_*c*_ emerges, where α represents the hydraulic conductance of the tissue segment and *P*_*c*_ the pressure required to initiate fracture. Using the conductance estimated from the slow-injection plateau (α≈0.03 mL/min kPa), the threshold injection rate for the s-CT sample is approximately 3.2 mL/min. Although the precise value depends on experimental conditions, this framework provides a mechanistic interpretation linking the macroscopic injection experiments with the microstructural processes revealed by the proposed experiment. Together, these findings support the interpretation of intramural injection as a form of hydraulic fracture occurring within a layered poroelastic biological material such as the aortic media.

## Limitations

This study has several limitations. First, the experimental model consisted of initially healthy porcine aortas, which do not reproduce the structural alterations commonly present in human aortic disease, including medial degeneration and adventitial remodeling. Second, although the injection platform enabled controlled biaxial loading and systematic variation of hydraulic parameters, the delamination was influenced by adventitial needle or catheter insertion and therefore does not fully replicate the native initiation pathways of spontaneous aortic dissection. Third, some experimental parameters were necessarily coupled, including needle size, penetration depth, and local tissue damage at insertion, which may have influenced the measured pressure thresholds and propagation patterns. Finally, the synchrotron phase-contrast microtomography experiments were performed on a single specimen with a limited field of view and sequential injection steps, such that the full spatial extent of delamination and the independent effects of the two injection regimes could not be completely resolved. Future studies using larger sample sizes, diseased or human tissues, and expanded high-resolution imaging protocols are needed to further generalize these findings.

## Clinical implications

Aging and pathological degeneration or remodeling of the aneurysmal wall are often associated with the accumulation of glycosaminoglycans and proteoglycans within the interlamellar spaces of the medial layer [32, 33]. Under certain conditions, these molecules can sequester water and lead to significant local swelling, contributing to the development of medial damage, most concerning when characterized by a pooling of mucoid material with associated local smooth muscle cell loss and degradation of elastic fibers. Such alterations may elevate stresses in the structural elements connecting adjacent elastic lamellae [34, 35]. When the tissue fails, these stresses can promote localized medial delamination, potentially occurring more readily and rapidly in the pre-elastic regime than observed in the present study. Although our experimental model does not capture effects of local glycosaminoglycan accumulation or the cyclic loading experienced in vivo, observations associated with pressurized flow of fluid within the media does have a clinical correlate.

## Conclusion

This study combined controlled biaxial mechanical loading, intramural fluid injection, camera tracking, synchrotron phase-contrast microtomography and histology to investigate biomechanical mechanisms governing intramural flow-induced damage within the aortic media. The results show that intramural injection induces a progressive transition from interlamellar fluid permeation to pressure-driven delamination, strongly modulated by axial stretch, injection rate, and local hydraulic boundary conditions. Higher axial stretches increased resistance to lamellar separation but promoted axial propagation of the permeation / delamination front, whereas higher injection rates favored abrupt medial delamination. Synchrotron imaging further revealed the microstructural processes underlying this transition, including local lamellar decohesion and the formation of fluid-filled delamination cavities. Together, these findings support the interpretation of aortic intramural injection as a hydraulic fracture process occurring within a layered poroelastic tissue and provide new insight into the mechanical factors that may regulate the initiation and propagation of aortic dissection.

## Supporting information

Cavinatp_et _al_Suppl

## Acknowledgments

This work was supported by the US National Institutes of Health (U01 HL142518), French Agence National de la Recherche (ANR-22-CPJ1-0090-01 and ANR-24-CE19-7688), and the Leducq Foundation (22CVD03 – erAADicate). We gratefully acknowledge Tim Weitkamp for his assistance with the s-CT experiments conducted on the ANATOMIX beamline, and Aurélien Baquié for providing the injection device and control software for the synchrotron phase-contrast microtomography setup.

## References

1. Nienaber, C. A., Clough, R. E., Sakalihasan, N., Suzuki, T., Gibbs, R., Mussa, F., & Pepper, J. (2016). Aortic dissection. Nature Reviews Disease Primers, 2(1), 16053. 10.1038/nrdp.2016.53

2. Fleischmann, D., Afifi, R. O., Casanegra, A. I., Elefteriades, J. A., Gleason, T. G., Hanneman, K., Roselli, E. E., Willemink, M. J., & Fischbein, M. P. (2022). Imaging and surveillance of chronic aortic dissection: A scientific statement from the American Heart Association. Circulation: Cardiovascular Imaging, 15(3). 10.1161/HCI.0000000000000075

3. Rylski, B., Schilling, O., & Czerny, M. (2023). Acute aortic dissection: evidence, uncertainties, and future therapies. European Heart Journal, 44(10), 813–821. 10.1093/eurheartj/ehac757

4. Brunet, J., Pierrat, B., & Badel, P. (2021). Review of Current Advances in the Mechanical Description and Quantification of Aortic Dissection Mechanisms. IEEE Reviews in Biomedical Engineering, 14, 240–255. 10.1109/RBME.2019.2950140

5. Sherifova, S., & Holzapfel, G. A. (2019). Biomechanics of aortic wall failure with a focus on dissection and aneurysm: A review. Acta Biomaterialia, 99, 1–17. 10.1016/j.actbio.2019.08.017

6. Nienaber, C. A., & Eagle, K. A. (2003). Aortic Dissection: New Frontiers in Diagnosis and Management. Circulation, 108(5), 628–635. 10.1161/01.CIR.0000087009.16755.E4

7. Krukenberg, E. (1920). Beiträge zur Frage des Aneurysma dissecans. Beitr Pathol Anat Allg Pathol, 67, 329–351.

8. Haverich, A., & Boyle, E. C. (2021). Aortic dissection is a disease of the vasa vasorum. JTCVS Open, 5, 30–32. 10.1016/j.xjon.2020.12.012

9. Nienaber, C. A., & Eagle, K. A. (2003). Aortic Dissection: New Frontiers in Diagnosis and Management. Circulation, 108(5), 628–635. 10.1161/01.CIR.0000087009.16755.E4

10. Brunet, J., Pierrat, B., Adrien, J., Maire, E., Lane, B. A., Curt, N., & Badel, P. (2023). In situ visualization of aortic dissection propagation in notched rabbit aorta using synchrotron X-ray tomography. Acta Biomaterialia, 155, 449–460. 10.1016/j.actbio.2022.10.060

11. Khoury, M. K., Stranz, A. R., & Liu, B. (2020). Pathophysiology of Aortic Aneurysms: Insights from Animal Studies. Cardiology and Cardiovascular Medicine, 4(4), 498–514. 10.26502/fccm.92920146

12. Roach, M. R., He, J. C., & Kratky, R. G. (1999). Tear propagation in isolated, pressurized porcine thoracic aortas. The Canadian Journal of Cardiology, 15(5), 569–575.

13. Roach, M. R., & Song, S. H. (1994). Variations in strength of the porcine aorta as a function of location. Clinical and Investigative Medicine. Medecine Clinique Et Experimentale, 17(4), 308–318.

14. Carson, M. W., & Roach, M. R. (1990). The strength of the aortic media and its role in the propagation of aortic dissection. Journal of Biomechanics, 23(6), 579–588. 10.1016/0021-9290(90)90050-D

15. Prokop, E. K., Palmer, R. F., & Wheat, M. W., Jr. (1970). Hydrodynamic Forces in Dissecting Aneurysms. Circulation Research. 10.1161/01.RES.27.1.121

16. van Baardwijk, C., & Roach, M. R. (1987). Factors in the propagation of aortic dissections in canine thoracic aortas. Journal of Biomechanics, 20(1), 67–73. 10.1016/0021-9290(87)90268-5

17. Tiessen, I. M., & Roach, M. R. (1993). Factors in the Initiation and Propagation of Aortic Dissections in Human Autopsy Aortas. Journal of Biomechanical Engineering, 115(1), 123–125. 10.1115/1.2895461

18. Tam, A. S. M., Catherine Sapp, M., & Roach, M. R. (1998). The effect of tear depth on the propagation of aortic dissections in isolated porcine thoracic aorta. Journal of Biomechanics, 31(7), 673–676. 10.1016/S0021-9290(98)00058-X

19. Humphrey, J. D., Kang, T., Sakarda, P., & Anjanappa, M. (1993). Computer-aided vascular experimentation: a new electromechanical test system. Annals of Biomedical Engineering, 21(1), 33–43. 10.1007/BF02368162

20. Ferruzzi, J., Vorp, D. A., & Humphrey, J. D. (2010). On constitutive descriptors of the biaxial mechanical behaviour of human abdominal aorta and aneurysms. Journal of The Royal Society Interface, 8(56), 435–450. 10.1098/rsif.2010.0299

21. Jacobs, K., Docter, D., de Smit, L., Korfage, H. A. M., Visser, S. C., Lobbezoo, F., & de Bakker, B. S. (2024). High resolution imaging of human development: shedding light on contrast agents. Neuroradiology, 66(9), 1481–1493. 10.1007/s00234-024-03413-z

22. Ban, E., Cavinato, C., & Humphrey, J. D. (2021). Differential propensity of dissection along the aorta. Biomechanics and Modeling in Mechanobiology, 20, 895–907. 10.1007/s10237-021-01418-8

23. Rolf-Pissarczyk, M., Schussnig, R., Fries, T.-P., Fleischmann, D., Elefteriades, J. A., Humphrey, J. D., & Holzapfel, G. A. (2025). Mechanisms of aortic dissection: From pathological changes to experimental and in silico models. Progress in Materials Science, 150, 101363. 10.1016/j.pmatsci.2024.101363

24. Yu, X., Suki, B., & Zhang, Y. (2020). Avalanches and power law behavior in aortic dissection propagation. Science Advances, 6(21). 10.1126/sciadv.aaz1173

25. Sugita, S., & Matsumoto, T. (2017). Multiphoton microscopy observations of 3D elastin and collagen fiber microstructure changes during pressurization in aortic media. Biomechanics and Modeling in Mechanobiology, 16(3), 763–773. 10.1007/s10237-016-0851-9

26. Amabili, M., Asgari, M., Breslavsky, I. D., Franchini, G., Giovanniello, F., & Holzapfel, G. A. (2021). Microstructural and mechanical characterization of the layers of human descending thoracic aortas. Acta Biomaterialia, 134, 401–421. 10.1016/j.actbio.2021.07.036

27. Cavinato, C., Murtada, S.-I., Rojas, A., & Humphrey, J. D. (2021). Evolving structure-function relations during aortic maturation and aging revealed by multiphoton microscopy. Mechanisms of Ageing and Development, 196, 111471. 10.1016/j.mad.2021.111471

28. Wang, R., Yu, X., & Zhang, Y. (2020). Mechanical and structural contributions of elastin and collagen fibers to interlamellar bonding in the arterial wall. Biomechanics and Modeling in Mechanobiology, 20, 93–106. 10.1007/s10237-020-01370-z

29. Stemper, B. D., Yoganandan, N., & Pintar, F. A. (2007). Mechanics of arterial subfailure with increasing loading rate. Journal of Biomechanics, 40(8), 1806–1812. 10.1016/j.jbiomech.2006.07.005

30. Bukac, M., Yotov, I., Zakerzadeh, R., & Zunino, P. (2015). Effects of Poroelasticity on Fluid-Structure Interaction in Arteries: a Computational Sensitivity Study. In A. Quarteroni (Ed.), Modeling the Heart and the Circulatory System (pp. 197–220). Cham: Springer International Publishing. 10.1007/978-3-319-05230-4_8

31. Leng, X., Zhou, B., Deng, X., Davis, L., Lessner, S. M., Sutton, M. A., & Shazly, T. (2018). Experimental and numerical studies of two arterial wall delamination modes. Journal of the Mechanical Behavior of Biomedical Materials, 77, 321–330. 10.1016/j.jmbbm.2017.09.025

32. Shen, Y. H., Lu, H. S., LeMaire, S. A., & Daugherty, A. (2019). Unfolding the Story of Proteoglycan Accumulation in Thoracic Aortic Aneurysm and Dissection. Arteriosclerosis, Thrombosis, and Vascular Biology, 39(10), 1899–1901. 10.1161/ATVBAHA.119.313279

33. Humphrey, J. D. (2013). Possible mechanical roles of glycosaminoglycans in thoracic aortic dissection and associations with dysregulated transforming growth factor-β. Journal of Vascular Research, 50(1), 1–10. 10.1159/000342436

34. Ahmadzadeh, H., Rausch, M. K., & Humphrey, J. D. (2019). Modeling lamellar disruption within the aortic wall using a particle-based approach. Scientific Reports, 9(1), 15320. 10.1038/s41598-019-51558-2

35. Liu, X., Ilseng, A., Prot, V., Skallerud, B. H., & Holzapfel, G. (2022). Swelling of interlamellar GAGs/PGs as an initiation mechanism for aortic dissection: Constitutive modeling and numerical simulations. Mechanics of Soft Materials, 4, 5. 10.1007/s42558-022-00043-4

