## Supplementary material for "Controlled intramural fluid injection to quantify propensity to thoracic aortic dissection": Cavinatp_et _al_Suppl

### Supplemental Figures

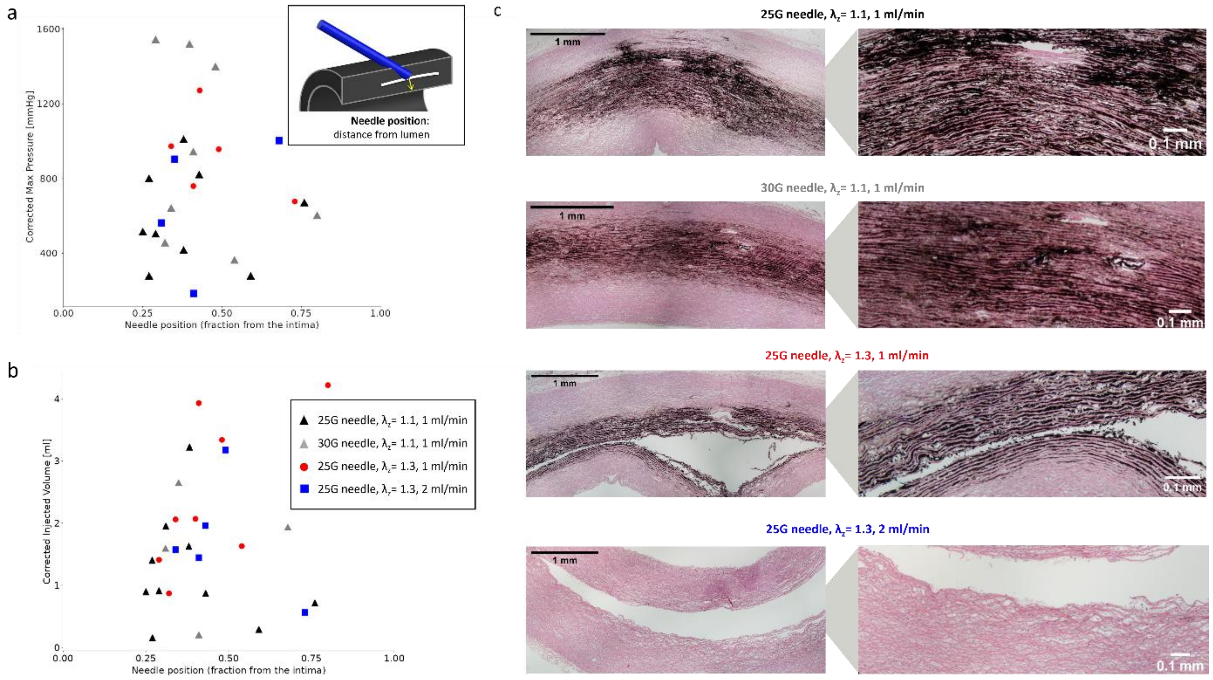


*Figure S1* ***Effects of needle penetration depth on injection pressure and delivered volume.*** *Scatter plots show the relationship between needle depth (measured from the intimal surface to the needle tip) and either (a) maximum injection pressure or (b) total injected volume at the pressure drop endpoint. Colors indicate experimental groups defined by needle gauge, axial stretch, and injection rate (black triangles: 25G needle, $\lambda_z$ = 1.1, 1 mL/min; grey triangles: 30G needle, $\lambda_z$ = 1.1, 1 mL/min; red circles: 25G needle, $\lambda_z$ = 1.3, 1 mL/min; blue squares: 25G needle, $\lambda_z$ = 1.3, 2 mL/min). The schematic illustrates the definition of penetration depth measured along the needle axis relative to the vessel wall curvature. (c) Representative histological cross-sections obtained at the axial site of fluid injection and stained with hematoxylin and eosin. Images were stitched together to reconstruct the arterial cross-section and analyzed to identify the radial position of the false lumen and quantify load-free wall thickness. The cases of injection at $\lambda_z$ = 1.1 presented multiple spots of localized lamellar decohesion in the area of fluid permeation, while all cases of injection at $\lambda_z$ = 1.3 consistently resulted in burst-like lamellar fracture.*

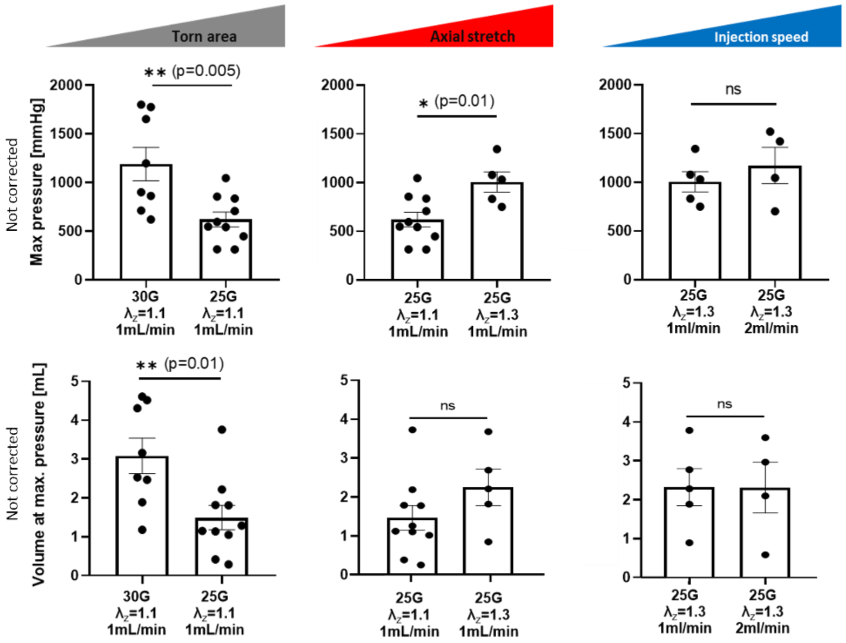


*Figure S2* ***Pressure-volume behavior during injection without corrections.*** *Bar plots summarizing mean maximum pressures (top) and corresponding injection volumes (bottom) for the four groups with no correction for tubing and needle effects (corrected data presented in Figure 3). Individual symbol types denote experimental conditions (axial stretch and injection rate) and the x-axis separates the groups according to needle gauge. Error bars represent SEM. Statistical significance, when present, is indicated by horizontal connectors above the bars.*

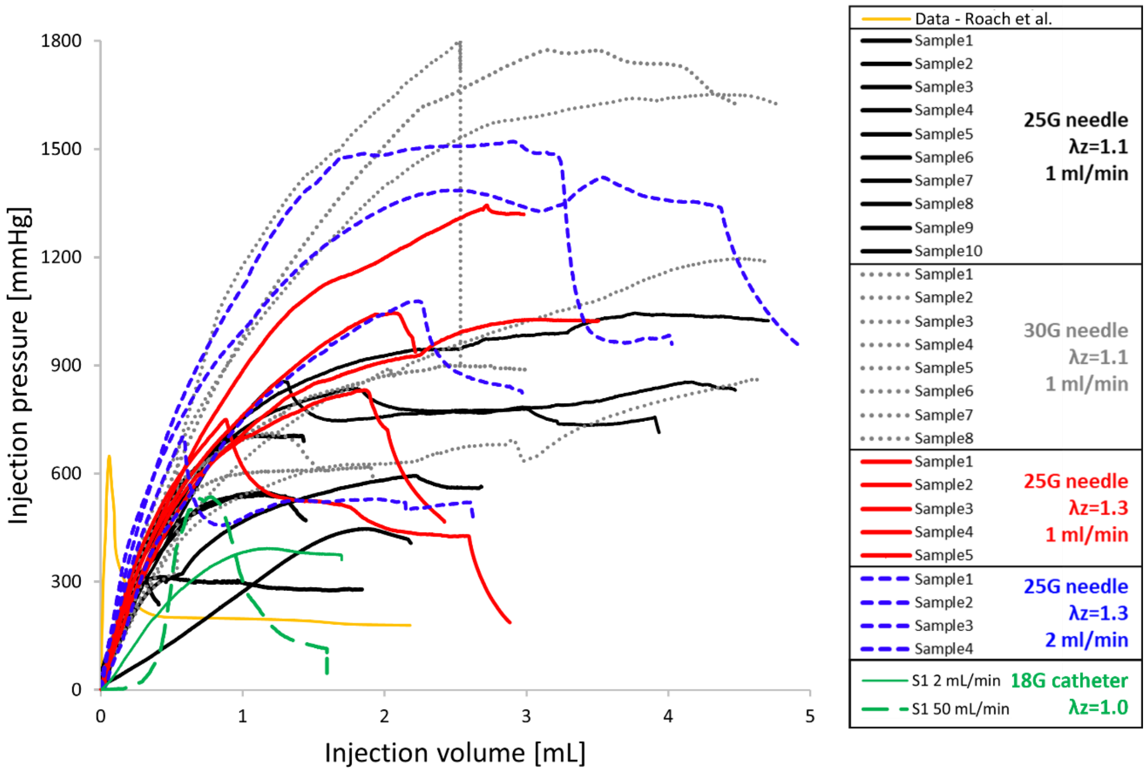


*Figure S3* ***Pressure-volume behavior during injection in comparison to data from Roach et al.*** [12]. *(a) Pressure–volume curves for porcine thoracic aortas injected using different experimental conditions as indicated in the legend: needle gauges (25G vs. 30G), fixed axial stretch during the injection ($\lambda_z$ = 1.1 or 1.3), and injection rates (1 or 2 mL/min), with luminal pressure fixed at 130 mmHg. Black solid lines: 25G needle, $\lambda_z$ = 1.1, 1 mL/min; dotted grey lines: 30G needle, $\lambda_z$ = 1.1, 1 mL/min; red solid lines: 25G needle, $\lambda_z$ = 1.3, 1 mL/min; blue dashed lines: 25G needle, $\lambda_z$ = 1.3, 2 mL/min. Green dashed line: reference data from one representative DTA sample from Roach et al. obtained using a 25G needle and saline containing India ink injected at a constant rate of 0.388 mL/min into the aorta pressurized with luminal pressure of 130 mmHg (axial stretch not reported); pressure represents the transmural pressure between the intramural bleb and the true lumen.*

### Supplemental Tables

*Table S1: Resulting parameters for the one-phase association model.*

| Conditions | YPlateau | K |
|---|---|---|
| 25G, 1 mL/min | 2924 | 0.353 |
| 30G, 1 mL/min | 2624 | 0.448 |
| 25G, 2 mL/min | 4021 | 0.210 |
| 30G, 2 mL/min | 2763 | 0.361 |

*Table S2: Pressure corrections, resulting from the Hagen-Poiseuille equation.*

| Conditions | 25G, 1mL/min | 30G, 1mL/min | 25G, 2mL/min | 30G, 2mL/min |
|---|---|---|---|---|
| Δp (mmHg) (tube+needle) | 36.285 | 259.239 | 72.570 | 518.477 |
